# The genome of the brackish-water malaria vector *Anopheles aquasalis*

**DOI:** 10.1101/2022.11.08.515629

**Authors:** Cesar C. P. Sepulveda, Rodrigo M. Alencar, Luiz Martinez-Villegas, Ana Cristina Bahia, Rosa A. Santana, Igor B. de Souza, Gigliola M. A. D’Elia, Ana Paula M. Duarte, Marcus V. G. de Lacerda, Wuelton M. Monteiro, Nágila F. Costa Secundino, Leonardo B. Koerich, Paulo F. P. Pimenta

## Abstract

*Anopheles aquasalis* is a primary malaria vector in coastal South America that grows in brackish waters of mangroves. Its importance has increased in recent years as it has been established as a model for parasite-vector studies for non-model *Plasmodium* species, such as *P. yoelli*. In this study, we present the complete genome of *An. aquasalis* and offer some insights into evolution and physiology. With a 162Mb and 12,446 coding proteins, the *An. aquasalis* genome is similar in size and gene content as other neotropical anophelines. 1,038 single-copy orthologs are present in *An. aquasalis* and all Diptera and it was possible to infer that *An. aquasalis* diverged from *An. darlingi* (the main malaria vector in inland South America) nearly 14 million years ago (mya). Ion transport and metabolism proteins is one the major gene families in *An. aquasalis* with 660 genes. Amongst these genes, important gene families relevant for osmosis control (e.g., aquaporins, vacuolar-ATPases, Na^+^/K^+^-ATPases and carbonic anhydrases) were identified in one-to-one orthologs with other anophelines. Evolutionary analysis suggests that all osmotic regulation genes are under strong purifying selection. We also observed low copy number variation in immunity-related genes (for which all classical pathways were described) and insecticide resistance genes. This is the third genome of a neotropical anopheline published so far. The data provided by this study may offer candidate genes for further studies on parasite-vector interactions and for studies on how brackish water anophelines deals with high fluctuation in water salinity.

**Significance Statement:** The brackish water mosquito *Anopheles aquasalis* is a primary malaria vector in coastal South America. Besides its peculiar ecological features (it is one of the few anopheline mosquitoes that survives high fluctuation of water salinity), *An. aquasalis* has gained relevance in recent years as a model for parasite-vector studies for non-model *Plasmodium* parasites. Still, the physiology and genetics of *An. aquasalis* are poorly understood. Here we present the genome of *An. aquasalis* with more than 12,000 annotated genes, offering insights in genome evolution, osmoregulation related, immunity, chemosensory and insecticide resistance genes. The data presented here will help to further advance the studies on *An. aquasalis* genetics and physiology to better understand parasite-vector interactions in non-model organisms.

## Introduction

Malaria is an acute febrile disease caused by *Plasmodium* sp. parasites that annually generates approximately 229 million cases and nearly half a million deaths worldwide (1). It is transmitted through the bite of a female mosquito of the *Anopheles* genus. Although this genus comprises 400 species, only 10% are recognized as vectors of the malaria parasite (2). Biological characteristics influenced by variations in the ability of these mosquitoes to transmit the parasite (e.g., molecular components of the immune response and intestinal physiology) are well studied and characterized in established models such as *An. gambiae* and *An. stephensi* (3–6). *Anopheles aquasalis* is a primary malaria vector in coastal South America, where the larvae develop in the brackish water of mangroves. *An. aquasalis* is among the few anophelines capable of surviving in severely changes in water salinity and rising sea levels due to climate changes has raised the concern of risk of disease transmission in coastal regions (7). However, only a few studies have addressed the question of how anopheline larvae deals with saline stress. Physiological studies have suggested that morphological changes in the localization of vacuolar-ATPases (V-ATPases) and K+/Na+ ATPase proteins are key for osmotic regulation in *An. albimanus* (8, 9). Other molecular and transcriptome studies also suggest that other genes, such as aquaporin and carbonic anhydrases, in conjunction with transcription modulation, are essential for mosquito survival in hyperosmotic or hypo-osmotic environments (10–12) Besides its peculiar ecological features, recent research recognized *An. aquasalis* as an important model for studying the interaction with human *Plasmodium* and murine models such as *P. yoelii nigeriensis* (13–15). Since then, it was possible to identify and functionally characterize the role of molecular components that are relevant during *Plasmodium’*s invasion of the mosquito midgut (13, 16–18).

However, despite its vectorial, ecological, and model importance, no genomic studies on *An. aquasalis* have been carried out so far. Genomic studies are fast and reliable methods for the genome exploration of medically important non-model insects (19, 20). Genomic and transcriptomic studies have given a better understanding of the genetic characteristics of more than 18 anopheline species, establishing the composition of conserved regions of genes, the identification of highly divergent genes, recognition of gene families, and the evolution of species-specific physiological or behavioral genetic variations (18, 21–23). Here we present the analysis of the genome of *An. aquasalis*. By identifying its coding genes, we were able to draw insights into genome evolution, genome structure, proteins relevant to osmotic regulation, genes relevant to vector-parasite and vector-host interactions, genes related to insecticide resistance, and more.

## Results

### Genome assembly and annotation of the *An. aquasalis* mosquito

The assembled *A. aquasalis* genome (BioProject PRJNA389759) has 162,944 Mb, distributed in 16,504 scaffolds (N50 14,431). 12,446 protein-coding genes were predicted in the genome (Data S1) and BUSCO analysis [92.1% of complete single-copy genes, 0.3% complete and duplicated, 2.2% fragmented and 5.4% missing genes] shows similar quality when compared to other mosquitoes (SI Appendix, Fig S1). The gene structure model generated with the MAKER program was evaluated according to the AED, with most gene structures supported by evidence, with 90% having a value between 0 and 0.5 of AED (SI Appendix, Fig. S2). Orthology analysis (Fig. 1B) suggests that 1,038 genes are single copy orthologs at Diptera level; 4,766 genes are multicopy genes with orthology to Diptera; 1,131 genes are present in *Drosophila melanogaster* (but not all anophelines); 1,999 genes are exclusive from mosquitoes; 308 genes are present in all anophelines; 1,016 genes are present in at least one other Anophelinae, 105 genes are exclusive from neotropical anophelines; 1,918 genes present orthology with other Diptera and 81 genes are of unknown origin. Based on the 1,038 single-copy orthologous genes, we reconstructed the evolutionary tree of *An. aquasalis* and other neotropical Anophelinae and calculated the divergence times for each branch (Fig. 1A). The phylogenetic tree suggests that *An. aquasalis* diverged from *An. darlingi* ∼14mya (Million Years Ago). South American anophelines diverged from *An. albimanus* ∼17mya and neotropical anophelines diverged from African anophelines ∼71mya.

**Fig. 1.**
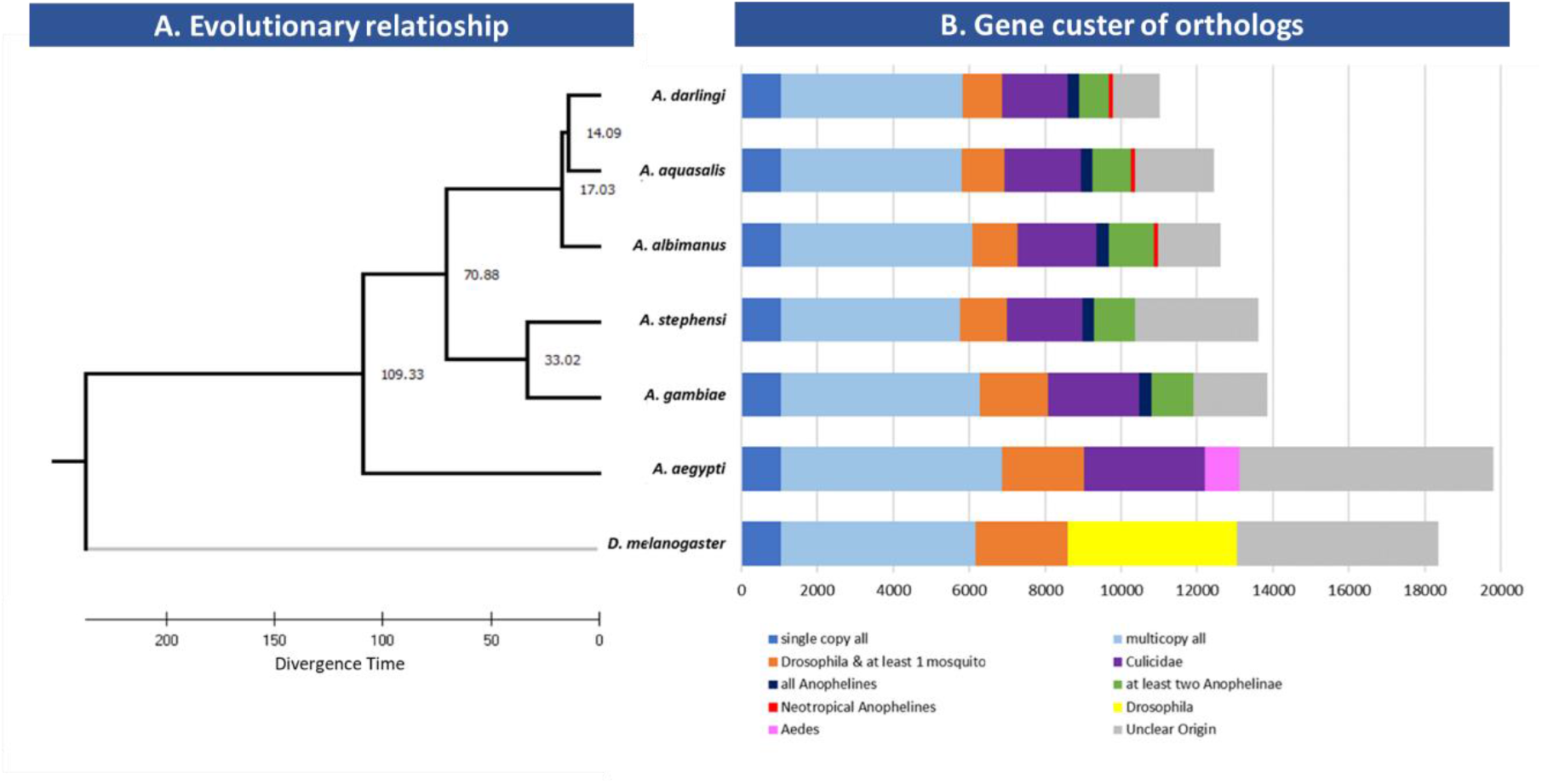
Evolutionary and orthology analysis of Neo Tropical anophelines. **A**. Phylogenetic tree was inferred based on 1038 single copy orthologs present in all taxa. Divergence time was calculated using two calibration constrains. **B**. Graphic bars show the orthology of annotated genes in each taxon. Single copy all: single copy orthologs present in all taxa; multicopy all: multicopy orthologs present in all taxa; *Drosophila &* at least 1 mosquito: orthologs present in *Drosophila* melanogaster and at least one mosquito; *Drosophila*: orthologs present in *D. melanogaster* and other Diptera, but absent in mosquitoes; Culicidae: orthologs present in all mosquitoes; *Aedes*: orthologs present only in *Aedes aegypti*; all Anophelines: orthologs present in all anophelines; at least two Anophelinae: orthologs present in more than one anopheline (but not all); Neotropical Anophelines: orthologs present only in neotropical Anophelines; Unclear Origin: specific genes (no orthology) or with orthology with other Diptera.

Gene cluster analysis (SI Appendix, Fig. S3) of orthologous genes suggests that 3,154 genes are specific to *An. aquasalis* when compared to *An. darlingi, An. albimanus* and *An. gambiae*. A cluster of 7,012 orthologs are present in all anophelines, while 331 orthologs are present only in neo-tropical anophelines. The mean transcript size was of 4059,96 bp, while coding sequences presented a mean size of 420.88bp (SI Appendix, Fig. S4). 35,352 introns were identified in the genome of *An. aquasalis* with an average size of 666.45bp. Finally, the average gene size was predicted to be 3508.37bp (SI Appendix, Tab. S1). The composition of repetitive elements in the genome of *An. aquasalis* is 0.95% (SI Appendix, Tab. S2).

### Functional prediction of encoded genes from the *An. aquasalis* mosquito genome

Putative function was inferred to 65.9% (8,208) of the predicted protein-coding genes (Fig. 2; Dataset S1). From the genes with identified ontologies and putative functions, our analysis indicated that most terms corresponded to the category of cellular processes and signaling (29.2%), followed by metabolism with 23.2%, and information storage and processes (12.4%) (Fig. 2). Looking at the more specific classification, the most abundant classes of genes belonged to signal transduction mechanisms (877 genes); transcription and transcription factors (759 genes); amino acid transport and metabolism (730 genes); inorganic ion transport and metabolism (660 genes); post-translational modification, protein turnover, and chaperones (636 genes) and; cell wall/membrane/envelope biogenesis (635 genes). Altogether, the genes in these six functional classes encompass 51.7% of all genes with ascertained putative function. As expected, most annotated genes have unknown functions (4,242 genes: 34.1%). 98 genes were classified into the mobilome functional class (transposons and prophages).

**Fig. 2.**
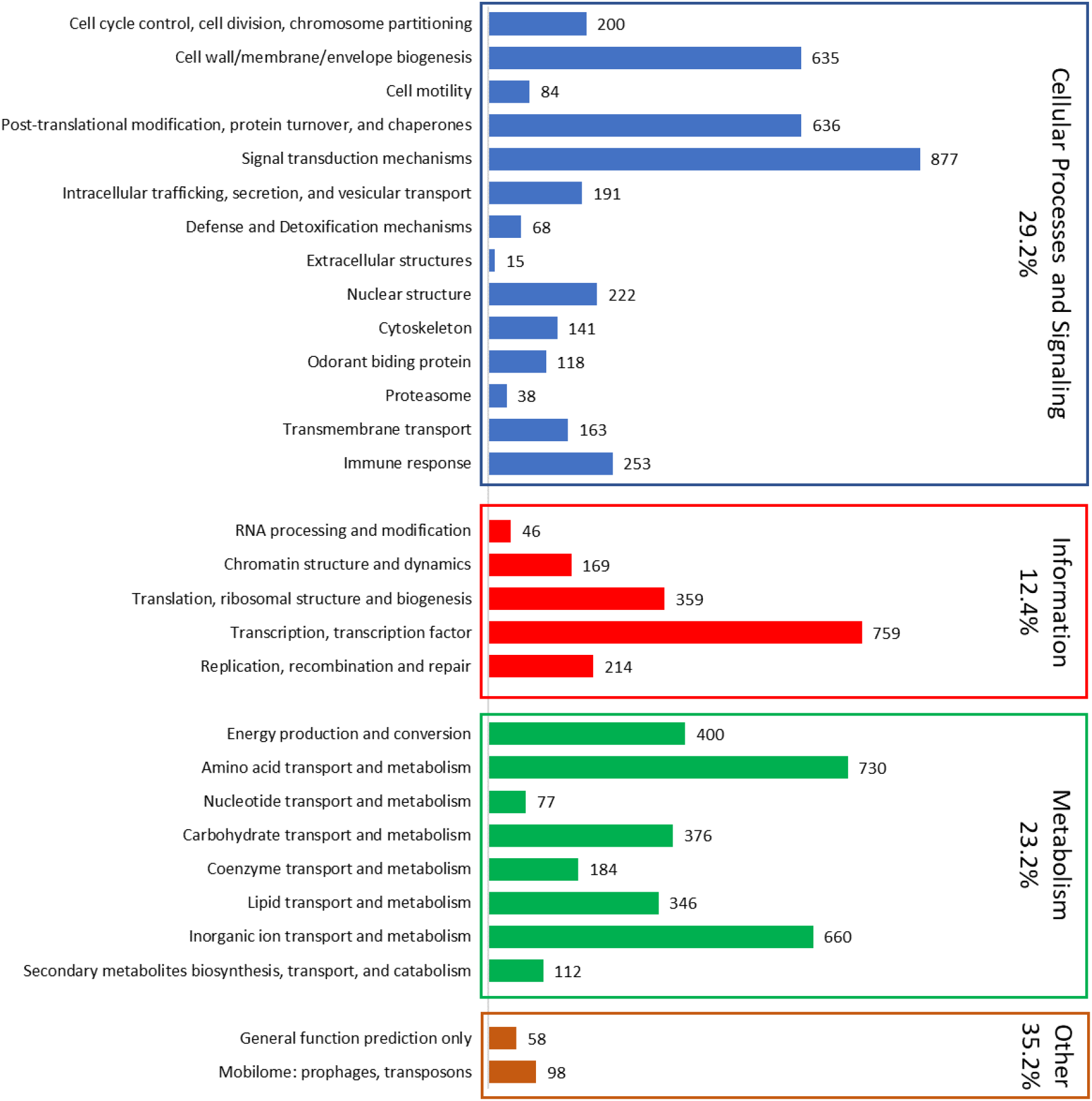
Summary of annotated genes in *An. aquasalis* genome. Bars represent the number of genes annotated into each functional class. Colors represent major functional groups and the percentage of genes in each category is represented. The percentage of genes belonging to the category of other genes (orange) includes genes with unknown functions (4,242 genes that were not represented by a bar in the figure due to scalability).

The analysis of the composition of domains with the InterproScan tool recognized among the main families of most representative proteins those that are composed of domains zinc finger C2H2-type (IPR013087) with 301 proteins, the Zinc finger, RING/FYVE/PHD-type (IPR013083) with 196 proteins and zinc finger, RING-type (IPR001841) with 101 proteins. Domains related to catalytic processes, such as protein kinase domains (IPR000719) with 212 proteins and serine proteases and trypsin domain (IPR001254) with 200 proteins also had a good representation. Other well-represented domains were those related to cell recognition processes, DNA repair, which are part of cell surface receptors and the immune response, such as immunoglobulin-like domain (IPR007110) with 168 proteins, Leucine-rich repeat, typical subtype (IPR003591) with 122 proteins, Leucine-rich repeat (IPR001611), RNA recognition motif domain (IPR000504) with 120 proteins and Fibronectin type III (IPR003961) with 64 proteins. Finally, some groups of domains related to the structure of the cuticle were abundant, such as Insect cuticle protein (IPR000618) with 97 proteins and the Chitin binding domain (IPR002557) with 96 proteins (SI Appendix; Table S2). In the next session, we will discuss some groups of interest.

### Osmoregulation, ion metabolism and transport

*An. aquasalis* larvae live in brackish waters and osmoregulation is a key process for these anophelines. However, osmoregulation is a complex process involving ion transport and metabolism genes, water permeability and tissue modifications. We found 685 genes related to ion transport and metabolism in *An. aquasalis*, which is remarkably higher than other anophelines, in special neo-tropical anophelines (Fig. 3A). *An. aquasalis* genome presents a higher number of zinc ion binding, calcium ion binding, potassium channel activity, calcium ion transport and sodium channel activity proteins than *An. darlingi, An. albimanus* and *An. gambiae* (Fig. 3B).

**Fig. 3.**
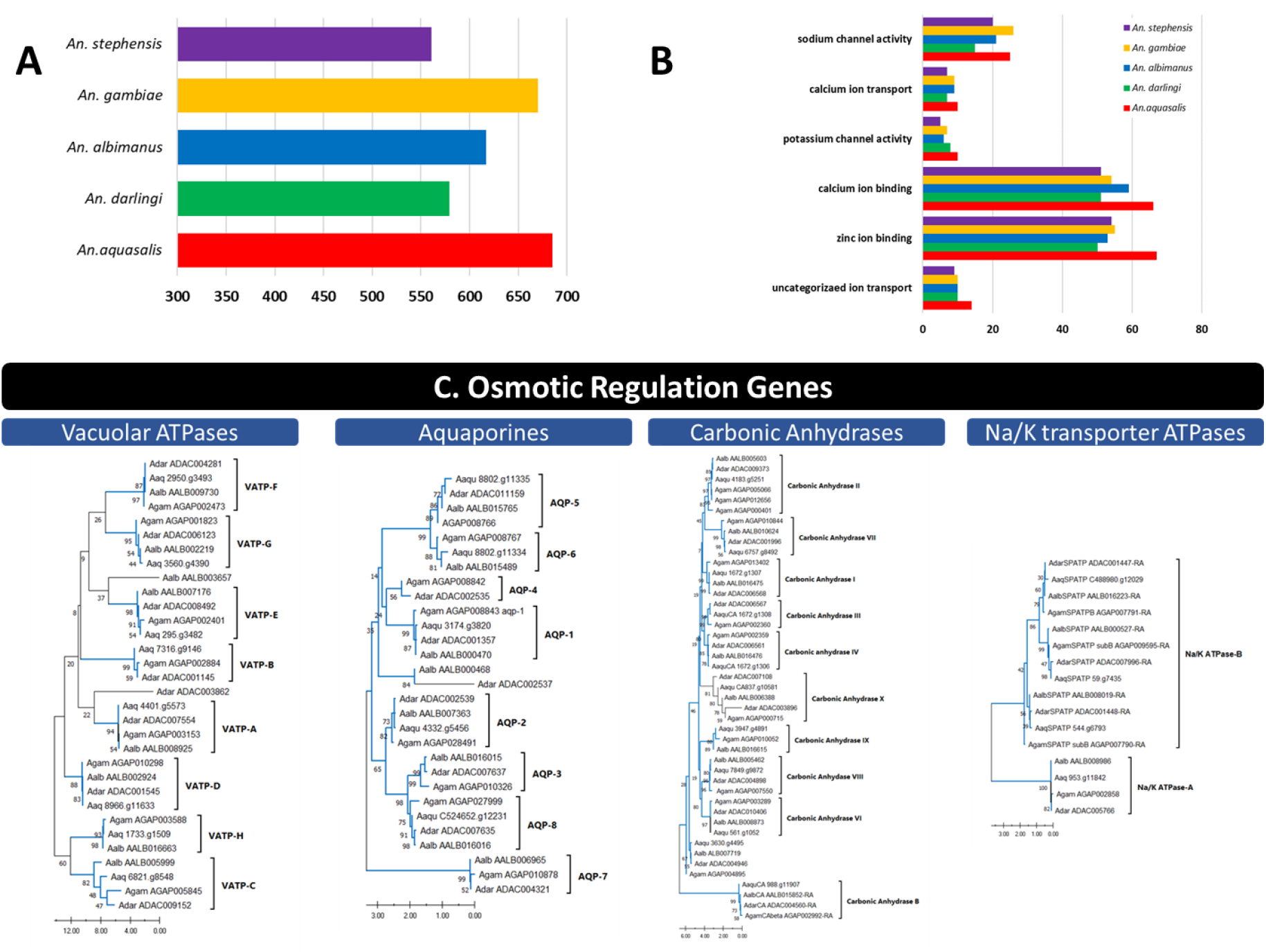
Ion metabolism and transport in *An. aquasalis*. Fig. 3A shows the number of genes related to ion metabolism and transport found in *An. aquasalis* in comparison to other anophelines. Fig. 3B shows the six families of genes related to ion metabolism and transport with higher abundance in *An. aquasalis* when compared to other anophelines. The osmotic regulation genes tab shows the evolutionary relationships of *An. aquasalis* (Aaq); *An. darlingi* (Adar); *An. albimanus* (Aalb) and *An. gambiae* (Agam) osmotic regulation genes. Trees were inferred by Neighbour Joining, using amino acid sequences, and bootstrapped (10,000 replicates). Blue branches indicate genes (and clades) under purifying selection tested by synonymous and nonsynonymous substitutions per site with p<0.05.

Studies on the tolerance of *An. albimanus* to saltwater also suggest the importance of V-type ATPase, carbonic anhydrase, and K+/Na+ ATPase proteins to osmoregulation (8–12), and studies in other insects have shown the role of aquaporins in such process. Orthology searches suggest that *An. aquasalis* has a similar, but slightly lower, number of osmoregulation genes (Fig. 3C) in comparison to neo-tropical anophelines. Evolutionary analyses and codon-based tests (Dataset S2) suggest that almost all osmotic regulation genes evolve under strong purifying selection (blue branches in Fig. 3C). Purifying selection was observed even between gene sub-families (e.g., all seven aquaporin gene subfamilies), except for vacuolar ATPases, in which evolutionary analysis suggests purifying selection within subfamilies but not between subfamilies (e.g., VATP-F and VATP-G).

### Immune response genes

The identification of molecular mechanisms of malaria vectors related to the decrease of *Plasmodium* sp. is of particular interest for understanding parasite-vector interactions. Hence, genes of the immune response system in *An. aquasalis* were discussed in depth in a companion paper (REFXXX) and here we will briefly present the most important findings and insights. We identified 278 immune related proteins, divided into 24 groups of families or signaling pathways, in the *An. aquasalis* genome. All genes from the classical signaling pathways (IMD, JAK/STAT, Toll and JNK) were identified with one-to-one orthologs to *An. darlingi*. Cascade modulators (e.g., serine proteases) accounted for 25.64% of identified immune response genes. Signaling pathways genes correspond to 13,46%. Other abundant families in the immune response group are FREPs with 29 genes (9.29%), autophagy process with 20 genes (6.41%), leucine-rich repeats (20; 6.41%) and C type lectins (13; 4.16%). In general, *An. aquasalis* has a similar number of immune response genes compared with Neo-tropical anophelines (*An. darlingi* 294; *An. albimanus* 304); but significantly lower number of genes when compared to *An. gambiae* (410).

### Chemosensory system

Anophelines use a series of chemosensory proteins to perceive its environments, such as the identification of hosts and oviposition sites. Chemosensory genes are generally classified as chemosensory proteins (CSPs) and odorant binding proteins (OBPs) and can be divided into three families of significant importance: odorant receptors (ORs), gustatory receptors (GRs) and ionotropic receptors (IRs). Our analysis found 44 ORs, 32 GRs and 15 IRs in the *An. Aquasalis* genome, which is a lower number than all anophelines so far, including *An. darlingi* with 57 ORs and 56 GRs. Despite the low conservation of OBPs amino acid sequences, we identified 6 conserved cysteine residues characteristic of this gene family. All *An. aquasalis* OBPs were also classified into Classic, Atypical, and Plus-C subfamilies according to their homology to *An. gambiae* sequences and phylogeny (SI Appendix, Fig. S5-S6, Table S4). Motif analysis identified eight conserved motifs in OBPs, with conserved cysteine residues in four of them (SI Appendix, Fig. S7)

### Insecticide resistance and detoxification

133 genes were identified related to metabolic detoxification, 73 (54%) from the P450s family (Table 1), 25 GST and 36 Carboxylesterases (SI Appendix, Tab. S5-S6). Evolutionary analyses with *An. gambiae* P450 genes (SI Appendix, Fig. S8-S10) allowed us to classify *An. aquasalis* P450 into four classical clans: CYP2 (8 genes), CYP3 (32 genes), CYP4 (25 genes) and mitochondrial CYP (8 genes). These numbers were similar to genes found in the genome of *An. darlingi*, however, *An. albimanus* and *An. gambiae* mosquitoes presented a greater number of genes.

**Table 1.** *An. aquasalis* Cytochrome P450 gene sub-families and comparison with *An. darlingi, An. albimanus* and *An. gambiae*.

|  | *An. aquasalis* | *An. darlingi* | *An. albimanus* | *An. gambiae* |
| --- | --- | --- | --- | --- |
| CYP2 | 8 | 10 | 10 | 11 |
| CYP3 | 32 | 32 | 35 | 39 |
| CYP4 | 25 | 22 | 36 | 45 |
| Mitochondrial CYP | 8 | 8 | 8 | 9 |
| **Total** | **73** | **72** | **89** | **104** |

Evolutionary analyses (SI Appendix, Fig. S5-S9) identified three *An. aquasalis* genes orthologous to *An. gambiae* genes that are related to insecticide resistance: 1921.g1856 is orthologous to AGAP002862 (CYP6AA1); XXX related to AGAPXXXX (CYP6M3) – both from the CYP3 family; and C559148.g12387 which is orthologous to AGAP001861-PA (CYP4H14) – from the CYP4 family. We also identified two losses of genes in the CYP2 family. The orthologous gene of CYP350 (AGAP005660, AALB015553 and ADAC003150), and the orthologous gene of CYP11179 (AGAP003065, AALB015657, ADAC007012). Finally, the *An. gambiae* mitochondrial CYP12F2 (AGAP008020-PA) and CYP12F3 (AGAP008019-PA) genes were not found in the genome of *An. aquasalis*.

Regarding Glutathione-S-transferases, genes were classified into seven classes: Delta, Epsilon, Omega, Sigma, Theta, Zeta and Unclassified (SI Appendix, Tab. S5). The number of genes of the main classes Delta and Epsilon did not have many alterations among the mosquitoes. Cholinesterases (CCEs) were classified into eight subfamilies: α-esterase, β-esterase, juvenile hormone esterase, acetylcholinesterase, gliotactin, glutactin, neurotactin and neuroligin (SI Appendix, Tab. S6). Among the identified CCEs, 36% belong to the α-esterase subfamily, the subfamily with the highest number of CCEs among anophelines (30%-36%).

## Discussion

The sequencing and annotation of anopheline genomes allowed the identification of structural, functional, and evolutionary differences of these mosquitoes. A task that started with *An. gambiae* at the beginning of this century and has so far studied more than 15 species (23–25). Five species of anophelines are responsible for most malaria transmission in the Americas. Two of them, *An. darlingi* and *An. albimanus* have already been sequenced. Here, we present the genome of the most important malaria coastal vector in the Americas, the *An. aquasalis*. Besides its importance as a vector of malaria parasites, *An. aquasalis* has a remarkable ecological feature since its larvae grow in the brackish water of mangroves. Hence, its genomic sequences could reveal genes and unique adaptations that deal with salted water, in contrast to other anophelines. Anophelines have genomes ranging from 134.7Mb (*An. darlingi*) to 375.8Mb (*An*. sinensis) – with a median size of 224.3Mb +-50,3Mb (23). The number of genes range from 10,457 (*An. darlingi*) to 16,149 (*An. melas*) with a mean of 13,162 genes +-1,380 (23). Our results suggest that the size of the genome (162.9 Mb) and several coding proteins (12,446) of *An. aquasalis* is relatively similar to other anophelines. We found a core of 1,038 single-copy genes with orthologs in all neotropical anophelines, *An. gambiae, An. stephensis* and *D. melanogaster*. Based on these single-copy orthologs, we could reconstruct the evolutionary history of *An. aquasalis* and estimate its divergence time from other anophelines. Our data suggest that *An. aquasalis* diverged from *An. darlingi* approximately 14 mya, and that neotropical anophelines diverged from old-world anophelines ∼70 mya. Previous works based on mitochondrial DNA, estimated the diverging time of neotropical anophelines to <u>African</u> anophelines between 79mya to 83.23 mya (26, 27). The work of Martinez-Villegas (26) was the first to estimate the *An. aquasalis* divergence time from other species and found the divergence of *An. aquasalis* to *An. darlingi* to be ∼39mya. On the other hand, the 16 *Anopheles* genomes manuscript suggests the divergence time of neotropical anophelines from African anophelines to be ∼100mya (23). However, the authors do not explain how they found this number or even if they calculated the divergence times. Hence, this manuscript is the first to calculate the divergence times based on a set of over a thousand nuclear genes, suggesting not only that *An. aquasalis* is more related to *An. darlingi* than to *An. albimanus*, but both species have a much earlier divergence than previously found.

Insects have developed several mechanisms to live in saline environments, from regulation of ion transport and metabolism genes to morphological modifications of the rectum. We found 660 proteins related to ion transport and metabolism, and our data suggest that *An. aquasalis* has a higher number of such proteins than neotropical anophelines and *An. gambiae*. Amongst the most abundant gene families related to ion transport, we identified many sodium channels (25 genes), potassium channels (10 genes) and calcium binding (66 genes) and transport (10 genes) proteins. *An. albimanus* is another anopheline with tolerance to saline water and presented a lower number of such genes (21 sodium channels; 6 potassium channels; 59 calcium binding and 9 calcium transports). However, the number of genes might not be the most important correlation. Physiological studies demonstrated that larvae of *An. albimanus*, when exposed to gradual changes in saline water, specialized non-dorsal anterior rectal (non-DAR) cells undergo changes in the localization of V-type ATPases and K+/Na+ ATPases proteins, allowing the production of super osmotic urine and disrupting the ion reabsorption system in non-DAR cells (8, 9). Other studies in mosquitoes have also suggested the importance of Carbonic Anhydrase (CA) and aquaporins to osmoregulation in saline environments (10, 12). We found that *An. aquasalis* has the number of such genes when compared to *An. albimanus*. Therefore, the ability of *An. aquasalis* to survive in saline environments does not seem to be related to an excess of copies of osmoregulation genes. We also hypothesized if the ability of *An. aquasalis* to live in saline water could be due to amino acid changes in such proteins (positive selection). However, as expected, our analysis suggests that all osmotic regulation genes are under strong purifying selection, within and between orthologous groups in each protein family (the exception are the V-type ATPases in which each ortholog groups seems to be evolving independently). Transcriptome studies in *An. merus* (another anopheline with high tolerance to saline environments) (11) revealed several changes in gene expression upon salinity stress, raising a few candidates for further functional studies. Therefore, further transcriptome studies on *An. aquasalis* may reveal significant changes in gene expression and indicate candidates for functional studies. *An. aquasalis* is one of the major malaria vectors in South America, and studies in parasite-vector and vector-host interactions have been the focus of many researchers. For the plasmodium to infect the anopheline, the parasite must be able to overcome the insect’s immune system. The *An. aquasalis* genome revealed all genes from classical immune pathways, in a one-to-one ortholog with *An. darlingi*. However, our study suggests that *An. aquasalis* (as *An. darlingi* and *An. albimanus*) has a lower number of immune-related genes than Old World anophelines.

In general, the family groups related to reactive oxygen species production control processes, and components regulating the expression of effectors of the immune response or signaling pathways were well conserved, with groupings of orthologs 1:1 for the four species. They are functionally relevant genes for the maintenance of the homeostasis of the organism, and in the case of signaling pathways such as the Toll pathway, they help in embryonic development processes, being constitutive for these insects (28–30). On the other hand, the marked differences in the number of copies were in groups related to the recognition of molecular patterns of microorganisms, with sharp differences in the FREP and MLD proteins, especially with species-specific expansions in *An. gambiae*. A phenomenon possibly induced by the microbiota of each species or by metabolic or sensory needs of each organism as in the case of MLD proteins (31–33).

Other families, such as PGRP or GNBP had few differences in the number of copies, with losses mainly in American anophelines. These proteins activate signaling pathways, and some have been found as regulatory factors for other members of the same family (34, 35). The regulatory role of both families allowed a few variations to be maintained during evolution in American anophelines and An. gambiae. Also, American mosquitoes suffered copy losses in cascade modulation proteins, mainly in serine proteases with CLIP domains, the most abundant family, and with gene expansions in An. gambiae. These gene families participate in activating signaling pathways and in the production of melanization components through proteolytic cascades after recognizing some pathogen (36, 37). It is recognized that specific sets of these proteins are organized for specific physiological and immune processes, sometimes with a redundant function that synergizes to increase the intensity of the response (38, 39). In this sense, it is speculated how exposure to specific pathogens shaped the set of serine proteases, serpins, and sometimes prophenoloxidase proteins in each species of mosquito (30, 37).

On the other spectrum, vector-host interactions, mosquitoes rely on a repertoire of chemosensory proteins to identify their hosts. Many studies demonstrated that CSPs and OBPs present rapid evolution, sometimes limiting the identification of orthologs even in close species. The 16 *Anopheles* genomes project suggested that most anophelines have ∼60 OR copies, while all anophelines of the gambiae complex gained ∼10 OR copies (23). On the other hand, GRs and IRs copy numbers remained stable in all species studies so far. Our data suggests that *An. aquasalis* has a much lower number of OR and GR genes than other anophelines (44 and 32 respectively). As rapid evolving genes, it is common that OBPs identification are underestimated in genomes, and this could be the reason to our findings. Neafsey and colleagues (23) tried to find a correlation in OBPs copy number variation (CNV) and host preference. However, transcriptome studies suggests that such differences are more likely due to functional divergence and regulation of gene expression (28, 29).

The increase of resistance to insecticides in insects’ vectors of diseases is of major concern for public health programs. Metabolic insecticide resistance is mediated by multi-copy gene families, such as cytochrome p450, glutathione S-transferases (GSTs) and carboxyl/cholinesterases. Despite its large numbers, several studies have shown the conservation of these gene families. We found 72 p450 from all relevant clades (CYP2, CYP3, CYP4 and mitochondrial CYP), a similar number of neotropical anophelines. The same is true for GSTs and carboxyl/cholinesterases. In most cases, we found a 1:1 ortholog to *An. gambiae*, including orthologs with genes related to insecticide resistance such as: CYP6AA1, CYP4H14 (resistance to pyrethroids), CYP6M2 (resistance to carbamates), CYP6M3 (resistance to organochlorines) (30, 31) and GSTE2, GSTE4, and GSTE6 (resistance do DDT, organochlorines and pyrethroids, respectively) (32, 33) We also observed a few gene duplications and losses and recent studies have suggested that CNV has a relevant roll in the rise of pyrethroid resistance (34, 35). We have no reports on the rise of insecticide resistance in *An. aquasalis* and only a few studies tried to address this issue (36, 37). Identifying orthologs of genes related to major insecticide resistance is relevant for future studies on insecticide metabolism and the evaluation of insecticide resistance in natural populations.

The data presented here opens new insights to *An. aquasalis* biology, neotropical anopheline evolutionary relationships and general anopheline evolution. Despite being a malaria vector in South America, the physiology of *An. aquasalis* is still poorly understood. Recent research elevated *An. aquasalis* as a significant model of vector-parasite interaction (13–15) and the identification of immunity and digestion-related genes are important for future research. Moreover, *An. aquasalis* is among the few anophelines capable of surviving drastic changes in water salinity and, with climate change and increased potential for saltwater invasion and salinization of inland waters (7), the study of the physiology of saltwater anophelines can be of great significance.

## Materials and Methods

### *Anopheles aquasalis* mosquito sampling and sequencing

The mosquito sample used for this work is from the colony established in the laboratory of Medical Entomology at FIOCRUZ-MG. Genomic DNA was purified from female pupae, using a Qiagen DNeasy Kit for blood and tissues. The library was prepared and sequenced as previously described by our group (38) and sequenced with Illumina HiSeq 2000 technology. The genome was assembled using SPADEs, also described by our group (38). The assembled genome is available under the accession number GCA_002846955.1.

### Databases and sequences used to predict the genome of *An. aquasalis*

As part of the evidence implemented for gene prediction, cDNAs generated from the transcriptome of (39) were downloaded from the GEO database with accession number GSE124997 on the NCBI website. In addition, proteins from four species of anophelines deposited in the VEuPathDB database were downloaded: *An. albimanus* (Anopheles-albimanus-STECLA_PEPTIDES_AalbS2.6), *An. darlingi* (Anopheles-darlingi-Coari_PEPTIDES_AdarC3.8.fa), *An. sinensis* (Anopheles-sinensis-China_PEPTIDES_AsinC2.2.fa) and *An. gambiae* (Anopheles-gambiae-PEST_PEPTIDES_AgamP4.12.fa) (40).

### Annotation and prediction of genes encoded by the genome of *An. aquasalis*

Simple repeats and complex repetitive elements in the genome of *An. aquasalis*, were predicted and masked with the RepeatMasker (http://repeatmasker.org) using the database of repetitive elements of the *An. gambiae* mosquito available on the VEuPathDB website with access on date 14/ 05/2020 (Anopheles-gambiae-PEST_REPEATS.lib) (41). The options for searching the elements of interest were –nolow to mask only the interspaced sequences. –norna to not mask the smallRNA genes. The file with the program’s predictions was obtained in gff format, with the – gff option (42).

### Structural prediction of the genome of *Anopheles aquasalis*

60,752 proteins from five anopheline species available in VEuPathDB and *An. aquasalis* transcripts (39) were used as a template for gene annotation. Initial prediction of genes was performed with MAKER program available online on the Galaxy server (43) in two rounds to create a draft of *An. aquasalis* putative genes. The MAKER archive and *An. aquasalis* ESTs were then used to train the AUGUSTUS program (44), and models generated by the training file was used for final annotation with AUGUSTUS using ESTs as evidence of transcription.

### Evaluation of the structural prediction of the *An. aquasalis* genome

The quality of the predictions was evaluated for each result obtained with the MAKER and Augustus programs using the BUSCO program, selecting the Diptera lineage and genome options on the Galaxy Australia server (43, 45). Also, the result obtained by the MAKER with the AED index was evaluated, establishing how much of the information used as evidence directly align with the *An. Aquasalis’* genome and this agreement between evidence and prediction must be equal to 90% of the generated alignments. The AED cumulative fraction of the annotations graph was generated using the AED_cdf_generator.pl script (46, 47).

### Functional prediction of the genome of the *An. aquasalis* mosquito

Protein sequences originating from the *An. aquasalis* genome gene model were used for functional prediction, gene ontology (GO) assignments, and functional descriptions of the *An. aquasalis* genome was generated through the Pannzer program pipeline, selecting those annotations with the ppv > 0.5 (48). In the case of protein domains and some gene ontologies (GO), they were annotated and searched with the InterproScan tool from the Galaxy Europe server, selecting the annotations as an e-score of 0.0001 (49). The REVIGO tool (http://revigo.irb.hr/) was used to identify some GO terms not identified by functional annotation programs and to reduce the redundancy of these terms. Finally, a homology search was made using the Blastp program, against the proteins of *Drosophila melanogaster* (Strain Berkeley) (UP000000803), *Culex quinquefasciatus* (Southern House mosquito Strain JHB) (UP000002320), *Aedes aegypti* (Yellow fever mosquito Strain LVP_AGWV) (UP000008820), *An. darlingi* (UP000000673), *An. albimanus* (New world malaria mosquito Strain STECLA/ALB19) (UP000069272) and *An. gambiae* (African malaria Pest strain) (UP000007062), choosing the sequences with a percentage identity >50 % and with e-value 0.0005 (50). The data generated by each tool were filtered using the programs, Excel and the online tool Google Collaboratory. To classify the terms of the genetic ontological obtained in the functional prediction, the R GO.db version 3.13.0 package was used, using the option “GOANCESTOR”. The functional prediction was complemented with orthology searches in OrthoDB v10.1 at Diptera Level(51)

### Orthology analysis

Orthology assignments were retrieved from OrthoDB v10.1 (51) Diptera-level orthologous groups (116 species) for the species detailed in Dataset S1. *An. aquasalis* protein-coding genes were mapped to OrthoDB v10.1 at the Diptera-level using *D. melanogaster* as an anchor. Mapping for each species was then merged to create the final orthologous groups including all mapped *An. aquasalis* proteins. The presence, absence, and copy-numbers of orthologs were assessed to partition genes from each of the 6 species into the distribution of categories shown in the bar chart (Fig. 1A) and classified as: 1) single copy orthologs present in all taxa; 2) multicopy orthologs present in all taxa; 3) orthologs present in *D melanogaster* and at least one mosquito; 4) orthologs present in *D. melanogaster* and other Diptera, but absent in mosquitoes; 5) orthologs present in all mosquitoes; 6) orthologs present only in *Aedes aegypti*; 7) orthologs present in all anophelines; 8) orthologs present in more than one anopheline (but not all); 9) orthologs present only in neotropical Anophelines; 10) specific genes (no orthology) or with orthology with other Diptera. Venn diagram was built with the VennDiagram package for R (52)

### Phylogenomics and gene evolutionary analysis

All evolutionary analyses were conducted in MEGA 11 (53). Protein coding sequences were translated to amino acids and aligned with Muscle. Phylogenetic trees were constructed using amino acid sequences, by Neighbor-Joining (for orthology analysis) (54) with 10,000 replicates and a complete deletion option. For species relationship, a time tree was inferred by applying the RelTime method (55, 56) to the phylogenetic tree. The time tree was computed using 2 calibration constraints based on data available at http://www.timetree.org/ (*A. aegypti* vs *An. gambiae* and *An. gambiae* vs. *An. darlingi*). Purifying and Positive selection hypotheses were tested by synonymous and nonsynonymous substitutions per site (dS and dN, respectively) in MEGA11. P values less than 0.05 are considered significant at the 5% level within and between orthologous groups for each gene family. The variance of the difference was computed using the analytical method. Analyses were conducted using the Nei-Gojobori method (57). All ambiguous positions were removed for each sequence pair (pairwise deletion option).

## Acknowledgments

This study was partially funded by the following Brazilian agencies: Foundation of the Institute Oswaldo Cruz (FIOCRUZ), Brazilian Council for Scientific and Technological Development (CNPq), Coordination for Improvement of Higher Education Personnel (CAPES), Minas Gerais and Amazon States Research Support Foundation (FAPEMIG and FAPEAM). This manuscript is a part of the PhD thesis developed by CCPS and RMA and supervised by PFPP and LBK. PFPP, NFCS, WMM and MVGL are productivity fellows of the Brazilian Council for Scientific and Technological Development (CNPq). The funders had no role in study design, data collection and analysis, decision to publish or preparation of the manuscript. All authors read and approved the final version of this article

