## Supplementary material for "The genome of the brackish-water malaria vector *Anopheles aquasalis*": Suplementary Information

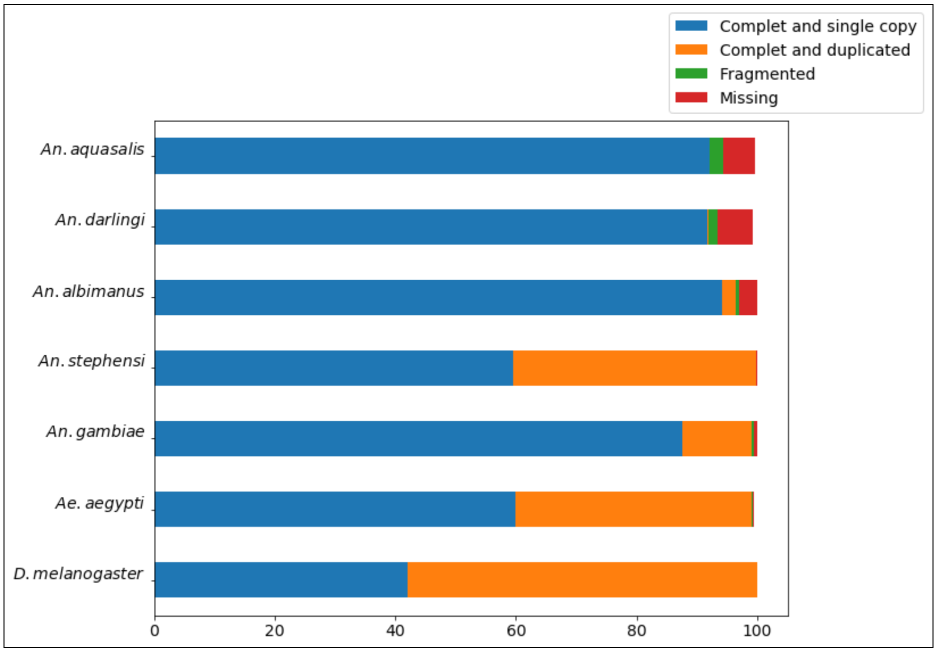


**Figure S1.** Comparison of the annotation integrity of the genomes of seven mosquito species evaluated with the BUSCO tool. Blue (complete and single-copy), Orange (Complete and duplicated), Green (fragmented), and red (missing).

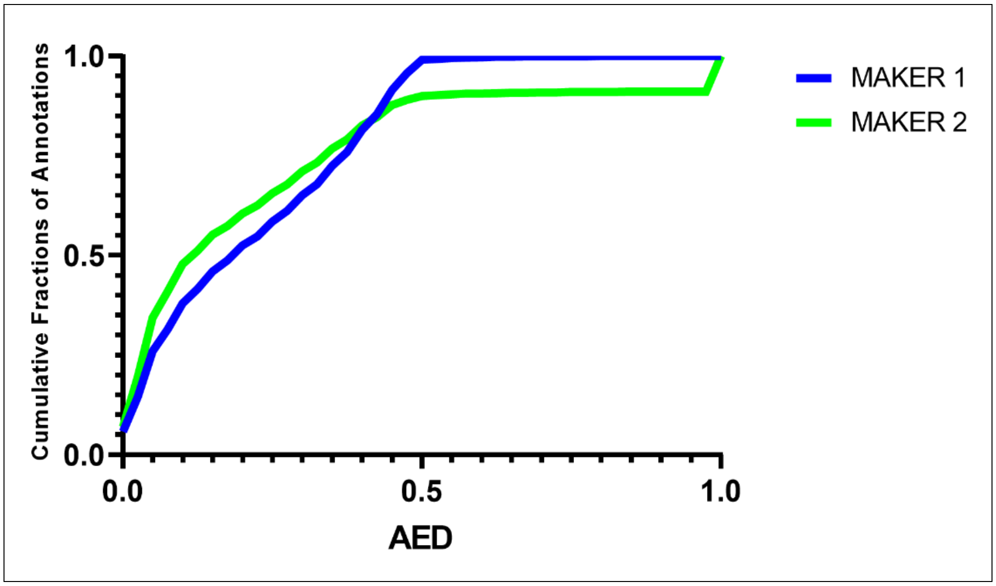


**Figure S2.** Evaluation of the annotation of the gene model generated by the MAKER program with the annotation edit distance index (AED). In blue is the result of the first annotation and in green is the result of the same index for the second annotation.

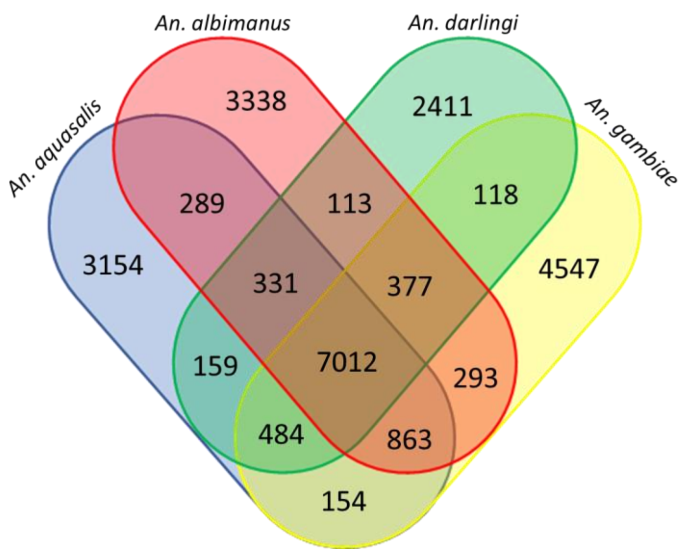


**Figure S3. Venn diagram of orthologs.** The Venn diagram shows the clustering of *An. aquasalis* genes (blue) identified by OrthoDB, in relation to four other anopheline species: *An. darlingi* (green); *An. albimanus* (red) and *An. gambiae* (yellow).

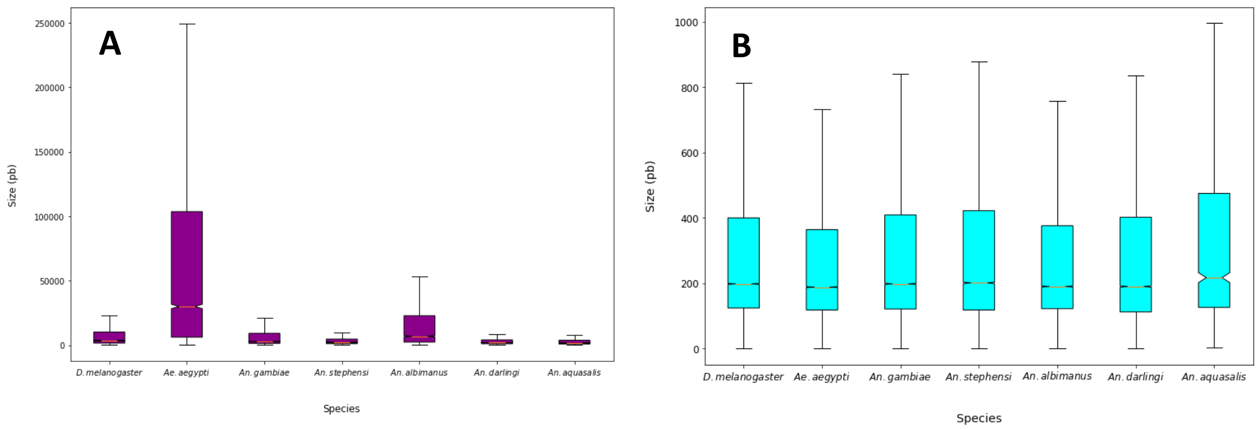


**Figure S4.** Structural comparison of genes among seven species of Diptera. A) Comparison of transcript size among seven species of Diptera. B) Comparison of the size of the coding region (CDS) among seven species of Diptera.

**Table S1.** General structural features of *An aquasalis* mosquito genes

| | **Max** | **Min** | **Mean** | **Median** |
|---|---|---|---|---|
| CDS | 13218 | 3 | 420,88 | 211 |
| Transcript | 63633 | 201 | 4059,96 | 1983 |
| Intron | 32585 | 55 | 684,4 | 84 |
| Gene | 63632 | 200 | 3508,37 | 1742,5 |

The composition of repetitive elements in the genome of *An. aquasalis* is 0.95%. Of these elements found in the genome of *An. aquasalis*, 0.40% were identified as retroelements and 0.21% as DNA transposons. The remaining 0.34% of the sequences were not categorized, so they were removed from masking. Among the transposable elements in *An. aquasalis*, most corresponded to the LINEs (30.55%), followed by the DNA transposable elements (22.3%) and finally the long terminal LTR repeats (11.22%) of the total finding with the tool by homology (Table S1).

**Table S2.** Composition of the DNA repetitive sequences of the *An. aquasalis* mosquito.

| Elements | Number of elements | Length | Percentage |
|---|---|---|---|
| **Retroelements** | 7290 | 503867 bp | **0.40 %** |
| **SINEs:** | 4 | 220 bp | 0.00 % |
| **Penelope** | 1 | 22 bp | 0.00 % |
| **LINEs:** | 5726 | 368137 bp | 0.29 % |
| **CRE/SLACS** | 0 | 0 bp | 0.00 % |
| **L2/CR1/Rex** | 80 | 7976 bp | 0.01 % |
| **R1/LOA/Jockey** | 388 | 30028 bp | 0.02 % |
| **R2/R4/NeSL** | 7 | 417 bp | 0.00 % |
| **RTE/Bov-B** | 44 | 7510 bp | 0.01 % |
| **L1/CIN4** | 119 | 5426 bp | 0.00 % |
| **LTR elements:** | 1560 | 135510 bp | 0.11 % |
| **BEL/Pao** | 173 | 32413 bp | 0.03 % |
| **Ty1/Copia** | 43 | 2895 bp | 0.00 % |
| **Gypsy/DIRS1** | 1341 | 99877 bp | 0.08 % |
| **Retroviral** | 0 | 0 bp | 0.00 % |
| **DNA transposons** | 3837 | 269843 bp | **0.21 %** |
| **hobo-Activator** | 2 | 183 bp | 0.00 % |
| **Tc1-IS630-Pogo** | 12 | 800 bp | 0.00 % |
| **En-Spm** | 0 | 0 bp | 0.00 % |
| **MuDR-IS905** | 0 | 0 bp | 0.00 % |
| **PiggyBac** | 1 | 40 bp | 0.00 % |
| **Tourist/Harbinger** | 5 | 533 bp | 0.00 % |
| **Other (Mirage,P-element, Transib)** | 54 | 4486 bp | 0.00 % |
| **Rolling-circles** | 29 | 4116 bp | 0.00 % |
| **Unclassified** | 4151 | 433291 bp | 0.34 % |
| **Total interspersed repeats** | | **1207001 bp** | **0.95 %** |

**Table S3.** Most abundant domains in the new world anopheline species *An. aquasalis*, *An. darlingi* and *An. albimanus.*

| | | Number of protein | | |
|---|---|---|---|---|
| InterPro ID | Description | *An. aquasalis* | *An. darlingi* | *An. albimanus* |
| IPR013087 | Zinc finger C2H2-type | 301 | 246 | 285 |
| IPR000719 | Protein kinase domain | 212 | 207 | 188 |
| IPR001254 | Serine proteases, trypsin domain | 200 | 195 | 188 |
| IPR013083 | Zinc finger, RING/FYVE/PHD-type | 196 | 182 | 191 |
| IPR007110 | Immunoglobulin-like domain | 168 | 142 | 175 |
| IPR001611 | Leucine-rich repeat | 163 | 144 | 150 |
| IPR001680 | WD40 repeat | 160 | 159 | 170 |
| IPR001314 | Peptidase S1A, chymotrypsin family | 154 | 157 | 144 |
| IPR003591 | Leucine-rich repeat, typical subtype | 122 | 103 | 112 |
| IPR000504 | RNA recognition motif domain | 120 | 115 | 119 |
| IPR003593 | AAA+ ATPase domain | 116 | 111 | 115 |
| IPR001841 | Zinc finger, RING-type | 101 | 96 | 99 |
| IPR000618 | Insect cuticle protein | 97 | 91 | 93 |
| IPR002557 | Chitin binding domain | 96 | 52 | 87 |
| IPR002110 | Ankyrin repeat | 92 | 85 | 99 |
| IPR001356 | Homeobox domain | 90 | 58 | 82 |
| IPR001128 | Cytochrome P450 | 87 | 93 | 99 |
| IPR000276 | G protein-coupled receptor, rhodopsin-like | 87 | 68 | 99 |
| IPR012934 | Zinc finger, AD-type | 78 | 76 | 89 |
| IPR001650 | Helicase, C-terminal | 77 | 75 | 80 |
| IPR000210 | BTB/POZ domain | 73 | 60 | 65 |
| IPR002048 | EF-hand domain | 73 | 70 | 77 |
| IPR001478 | PDZ domain | 72 | 61 | 70 |
| IPR001849 | Pleckstrin homology domain | 66 | 68 | 73 |
| IPR003961 | Fibronectin type III | 64 | 63 | 67 |
| IPR001806 | Small GTPase | 60 | 57 | 58 |
| IPR001452 | SH3 domain | 60 | 64 | 65 |
| IPR006170 | Pheromone/general odorant binding protein | 55 | 32 | 40 |
| IPR003439 | ABC transporter-like, ATP-binding domain | 54 | 55 | 48 |
| IPR011701 | Major facilitator superfamily | 53 | 56 | 56 |
| IPR001245 | Serine-threonine/tyrosine-protein kinase, catalytic domain | 49 | 54 | 35 |
| IPR001251 | CRAL-TRIO lipid binding domain | 48 | 49 | 37 |

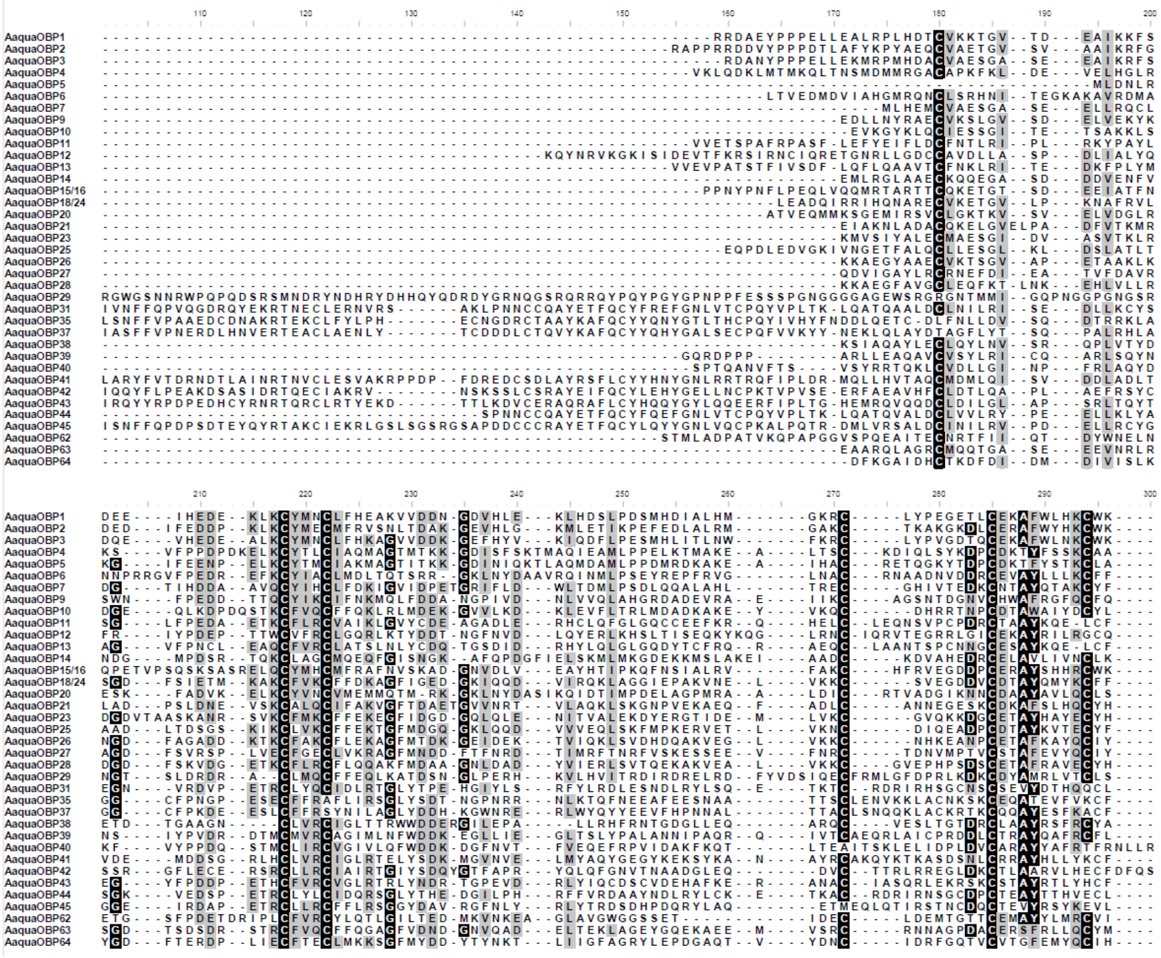

**Figure S5.** Multiple alignments of *An. aquasalis* OBPs to identify conserved cysteine residues.

**Table S4.** OBP genes found in the genome of the *An. aquasalis* mosquito

| Query | Aqua OBP | Subfamily | Size CDS | Size aa | Signal P start | Signal P end | CDD short name | CDD Superfamily | PFAM INTERPRO |
|---|---|---|---|---|---|---|---|---|---|
| 2978.g3569 | AaquaOBP43 | Atypical | 1023 | 340 | 1 | 23 | PBP_GOBP | cl11600 | PBP/GOBP family |
| 8779.g11276 | AaquaOBP9 | Classic | 420 | 139 | 1 | 17 | PBP_GOBP | cl11600 | PBP/GOBP family |
| 8493.g10819 | AaquaOBP63 | Classic | 357 | 118 | #N/A | #N/A | PBP_GOBP | cl11600 | PBP/GOBP family |
| 4149.g5178 | AaquaOBP31 | Atypical | 1275 | 424 | 1 | 22 | PBP_GOBP | cl11600 | PBP/GOBP family |
| 4149.g5179 | AaquaOBP45 | Atypical | 1362 | 453 | 1 | 26 | PBP_GOBP | cl11600 | PBP/GOBP family |
| 8493.g10820 | AaquaOBP26 | Classic | 396 | 131 | 1 | 18 | PBP_GOBP | cl11600 | PBP/GOBP family |
| 6912.g8608 | AaquaOBP41 | Atypical | 858 | 285 | 1 | 29 | PBP_GOBP | cl11600 | PBP/GOBP family |
| 1215.g336 | AaquaOBP39 | Atypical | 891 | 296 | 1 | 32 | PBP_GOBP | cl11600 | PBP/GOBP family |
| C493700.g12054 | AaquaOBP35 | Atypical | 831 | 276 | 1 | 20 | PBP_GOBP | cl11600 | PBP/GOBP family |
| 8493.g10822 | AaquaOBP18/24 | Classic | 471 | 156 | #N/A | #N/A | PBP_GOBP | cl11600 | PBP/GOBP family |
| 8493.g10816 | AaquaOBP28 | Classic | 417 | 138 | 1 | 20 | PBP_GOBP | cl11600 | PBP/GOBP family |
| 6459.g8135 | AaquaOBP15/16 | Classic | 516 | 171 | #N/A | #N/A | PBP_GOBP | cl11600 | PBP/GOBP family |
| 2191.g2289 | AaquaOBP42 | Atypical | 966 | 321 | 1 | 21 | PBP_GOBP | cl11600 | PBP/GOBP family |
| 8493.g10821 | AaquaOBP25 | Classic | 447 | 148 | 1 | 20 | PBP_GOBP | cl11600 | PBP/GOBP family |
| 1386.g768 | AaquaOBP20 | Classic | 483 | 160 | 1 | 39 | PBP_GOBP | cl11600 | #N/A |
| 8493.g10823 | AaquaOBP23 | Classic | 414 | 137 | 1 | 22 | PBP_GOBP | cl11600 | PBP/GOBP family |
| 3948.g4896 | AaquaOBP1 | Classic | 555 | 184 | 1 | 25 | PBP_GOBP | cl11600 | PBP/GOBP family |
| 8851.g11407 | AaquaOBP3 | Classic | 468 | 155 | 1 | 31 | PBP_GOBP | cl11600 | PBP/GOBP family |
| 1798.g1678 | AaquaOBP10 | Classic | 414 | 137 | 1 | 22 | PBP_GOBP | cl11600 | PBP/GOBP family |
| 2824.g3290 | AaquaOBP37 | Atypical | 897 | 298 | 1 | 22 | PBP_GOBP | cl11600 | PBP/GOBP family |
| 6459.g8134 | AaquaOBP2 | Classic | 498 | 165 | #N/A | #N/A | PBP_GOBP | cl11600 | PBP/GOBP family |
| 460.g5824 | AaquaOBP21 | Classic | 420 | 139 | 1 | 21 | PBP_GOBP | cl11600 | PBP/GOBP family |
| 1215.g334 | AaquaOBP12 | Classic | 741 | 246 | 1 | 22 | PBP_GOBP | cl11600 | PBP/GOBP family |
| 1863.g1758 | AaquaOBP4 | Classic | 543 | 180 | #N/A | #N/A | PBP_GOBP | cl11600 | PBP/GOBP family |
| 5301.g6641 | AaquaOBP6 | Classic | 486 | 161 | 1 | 27 | PBP_GOBP | cl11600 | PBP/GOBP family |
| 8150.g10277 | AaquaOBP7 | Classic | 339 | 112 | #N/A | #N/A | PBP_GOBP | cl11600 | PBP/GOBP family |
| 8973.g11644 | AaquaOBP14 | PlusC* | 432 | 143 | 1 | 20 | PBP_GOBP | cl11600 | PBP/GOBP family |
| 1215.g337 | AaquaOBP40 | Atypical | 1041 | 346 | 1 | 36 | PBP_GOBP | cl11600 | PBP/GOBP family |
| 397.g4962 | AaquaOBP5 | Classic | 291 | 96 | #N/A | #N/A | PBP_GOBP | cl11600 | PBP/GOBP family |
| 8493.g10817 | AaquaOBP64 | Classic | 426 | 141 | 1 | 28 | PBP_GOBP | cl11600 | PBP/GOBP family |
| 8493.g10818 | AaquaOBP27 | Classic | 414 | 137 | 1 | 25 | PBP_GOBP | cl11600 | PBP/GOBP family |
| 651.g8208 | AaquaOBP38 | Atypical | 1227 | 408 | 1 | 25 | PBP_GOBP | cl11600 | PBP/GOBP family |
| 1733.g1521 | AaquaOBP11 | Classic | 588 | 195 | 1 | 23 | PBP_GOBP | cl11600 | PBP/GOBP family |
| 122.g355 | AaquaOBP20_2 | PlusC* | 2256 | 751 | 1 | 39 | PBP_GOBP | cl11600 | #N/A |
| 1215.g335 | AaquaOBP13 | Classic | 573 | 190 | 1 | 21 | PBP_GOBP | cl11600 | PBP/GOBP family |
| 4149.g5177 | AaquaOBP44 | Atypical | 873 | 290 | #N/A | #N/A | PBP_GOBP | - | PBP/GOBP family |
| 157.g1071 | AaquaOBP157.g1071 | Classic | 837 | 278 | #N/A | #N/A | #N/A | #N/A | #N/A |
| 4433.g5610 | AaquaOBP62 | Classic | 501 | 166 | 1 | 26 | PBP_GOBP | - | PBP/GOBP family |
| 1180.g281 | AaquaOBP29 | Classic | 900 | 299 | 1 | 24 | PBP_GOBP | cl11600 | PBP/GOBP family |
| 5143.g6430 | AaquaOBP5143.g6430 | Classic | 948 | 315 | #N/A | #N/A | #N/A | #N/A | #N/A |
| 261.g3012 | AaquaOBP57 | PlusC | 504 | 167 | #N/A | #N/A | #N/A | #N/A | PBP/GOBP family |
| 6154.g7740 | AaquaOBP80 | PlusC | 618 | 205 | 1 | 29 | #N/A | #N/A | PBP/GOBP family |
| 3241.g3964 | AaquaOBP46 | PlusC | 624 | 207 | 1 | 24 | #N/A | #N/A | PBP/GOBP family |
| 3241.g3961 | AaquaOBP61 | PlusC | 603 | 200 | 1 | 19 | #N/A | #N/A | #N/A |
| 3241.g3960 | AaquaOBP60 | PlusC | 597 | 198 | 1 | 18 | #N/A | #N/A | #N/A |
| 3241.g3966 | AaquaOBP47 | PlusC | 582 | 193 | 1 | 21 | #N/A | #N/A | #N/A |
| 5805.g7304 | AaquaOBP58 | PlusC | 795 | 264 | 1 | 21 | #N/A | #N/A | #N/A |
| 3241.g3967 | AaquaOBP48 | PlusC | 591 | 196 | 1 | 25 | #N/A | #N/A | #N/A |
| 157.g1070 | AaquaOBP157.g1070 | Classic | 705 | 234 | #N/A | #N/A | #N/A | #N/A | #N/A |
| 3241.g3962 | AaquaOBP66 | Classic* | 525 | 174 | #N/A | #N/A | #N/A | #N/A | PBP/GOBP family |

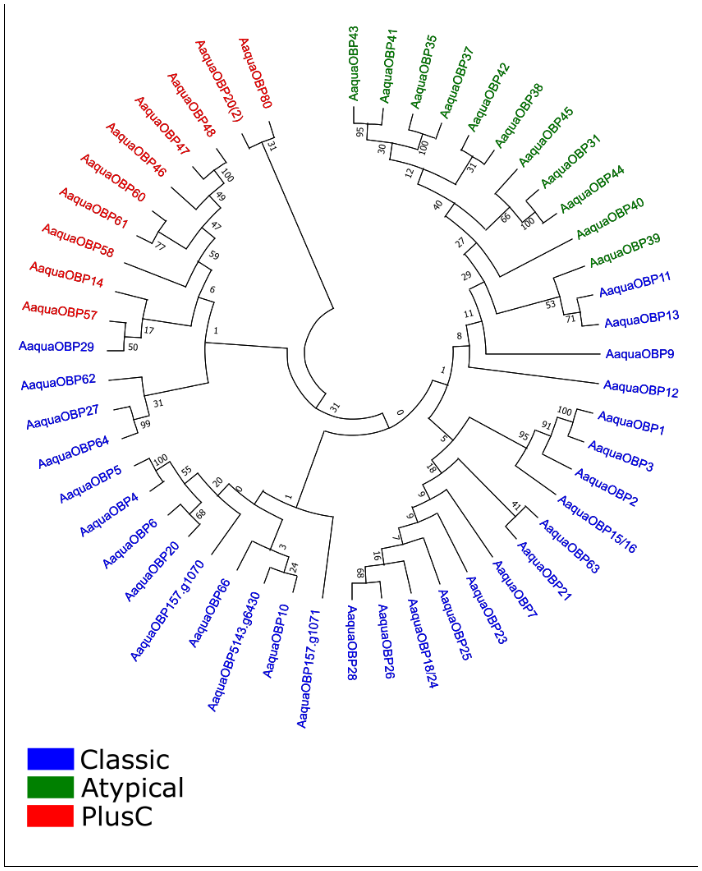


**Figure S6.** Neighbor-joining tree of OBPs amino acid sequences of *An. aquasalis*. Classification into Classic, Atypical and PlusC subfamilies

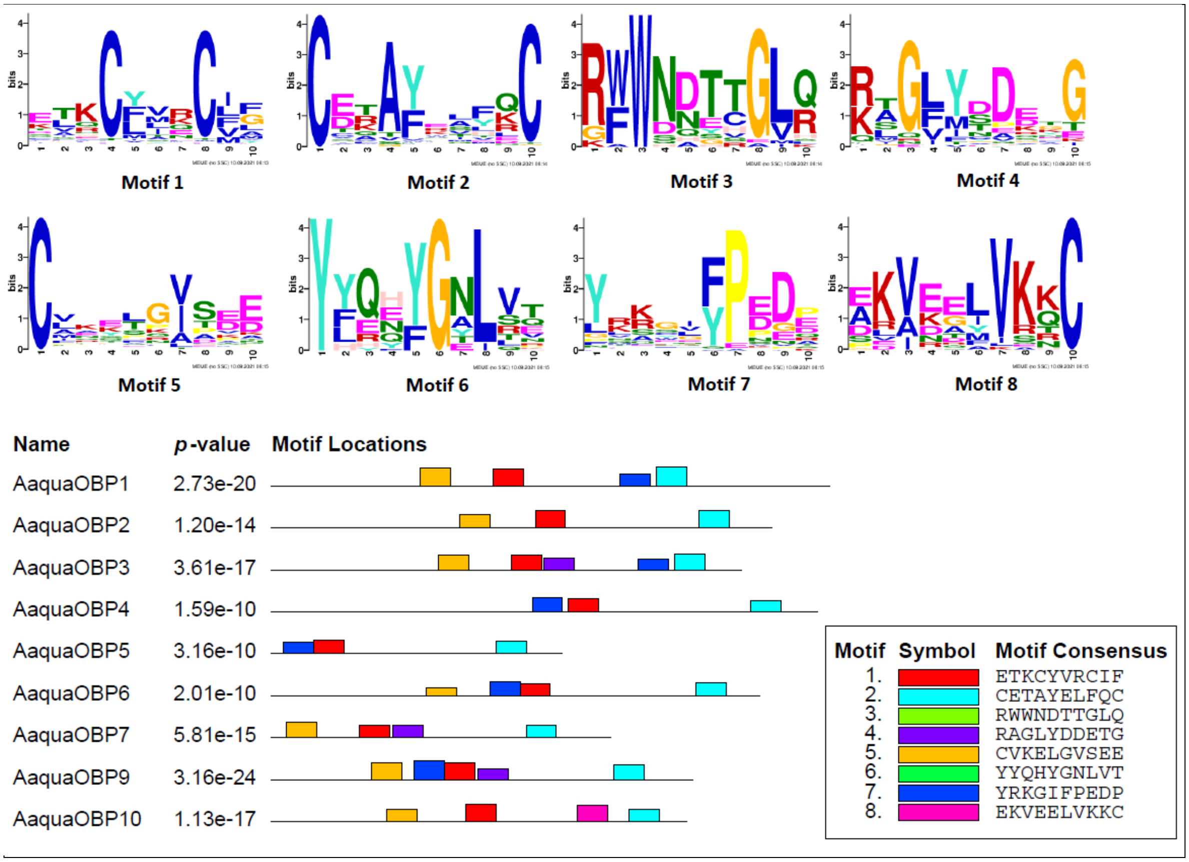


**Figure S7.** Analysis of OBP motifs from *An. aquasalis*, *An. gambiae*, *An. darlingi* and *An. albimanus.*

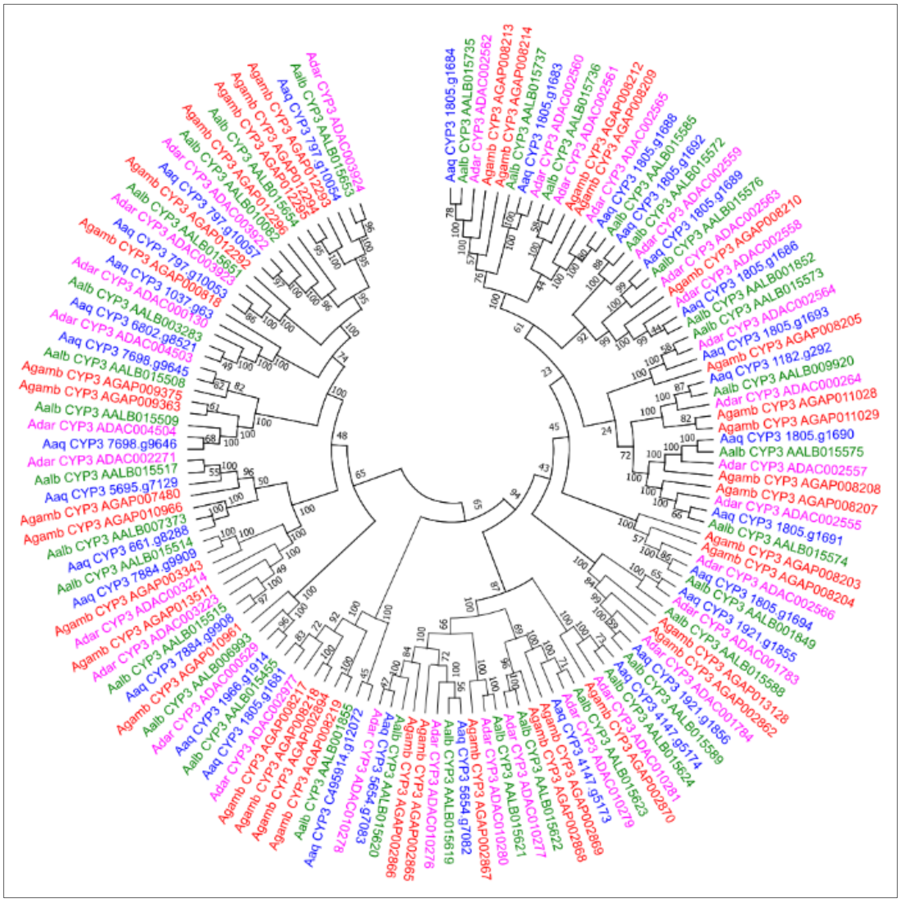

**Figure S8.** Neighbor-joining tree of CYP3 amino acid sequences of *An. aquasalis* (blue), *An. darlingi* (pink), *An. albimanus* (green) and *An. gambiae* (red) mosquitoes. Bootstrap values were calculated with 1000 replicates and their values were presented in each branch of the tree.

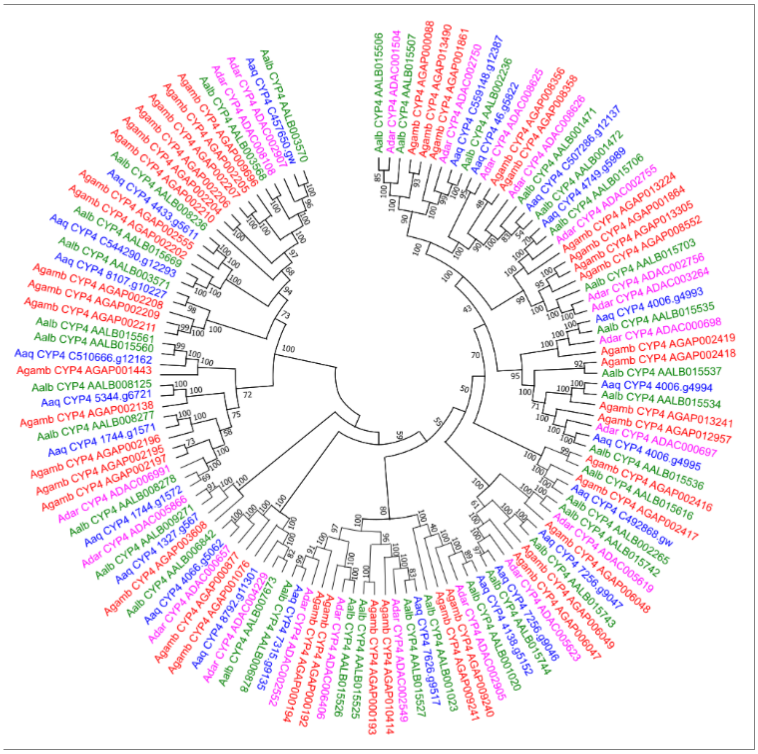

**Figure S9.** Neighbor-joining tree of CYP4 amino acid sequences of *An. aquasalis* (blue), *An. darlingi* (pink), *An. albimanus* (green) and *An. gambiae* (red) mosquitoes. Bootstrap values were calculated with 1000 replicates and their values were presented in each branch of the tree.

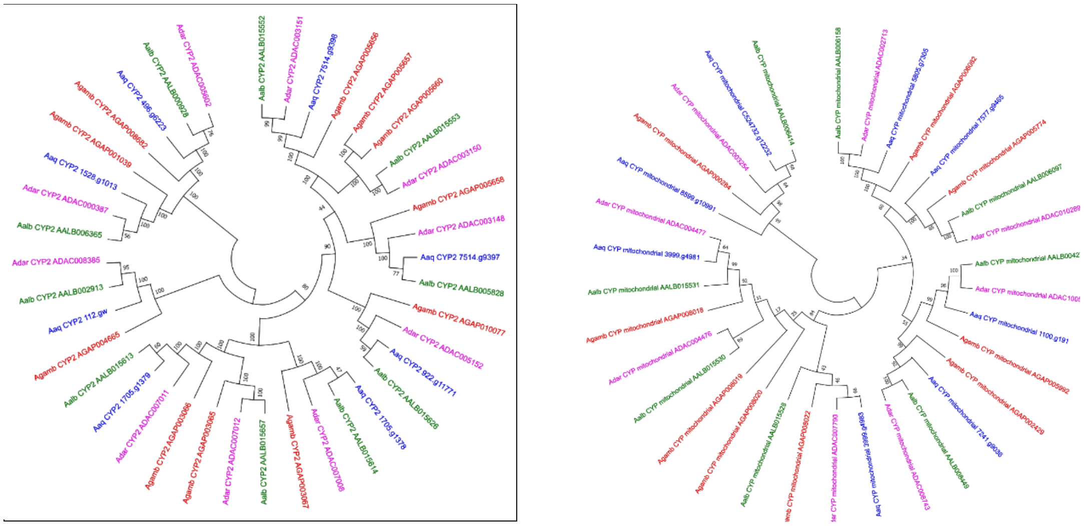

**Figure S10.** Neighbor-joining tree of CYP2 (A) and Mitochondrial CYP (B) amino acid sequences of *An. aquasalis* (blue), *An. darlingi* (pink), *An. albimanus* (green) and *An. gambiae* (red) mosquitoes. Bootstrap values were calculated with 1000 replicates and their values were presented in each branch of the tree.

**Table S5.** *An. aquasalis* Glutathione-S-transferases gene sub-families and comparison with *An. darlingi*, *An. albimanus* and *An. gambiae*.

| | *An. aquasalis* | *An. darlingi* | *An. albimanus* | *An. gambiae* |
|---|---|---|---|---|
| Delta | 11 | 11 | 10 | 11 |
| Epsilon | 5 | 6 | 6 | 6 |
| Omega | 1 | 1 | 1 | 1 |
| Sigma | 2 | 2 | 1 | 2 |
| Theta | 2 | 2 | 2 | 2 |
| Zeta | 1 | 1 | 1 | 1 |
| Unclassified | 3 | 3 | 3 | 3 |
| **Total** | **25** | **26** | **24** | **26** |

**Table S6.** *An. aquasalis* Cholinesterases gene sub-families and comparison with *An. darlingi*, *An. albimanus* and *An. gambiae*.

| | *An. aquasalis* | *An. darlingi* | *An. albimanus* | *An. gambiae* |
|---|---|---|---|---|
| α-esterase | 13 | 14 | 11 | 14 |
| β-esterase | 2 | 3 | 1 | 5 |
| Juvenile hormone | 3 | 4 | 2 | 5 |
| Acetylcholinesterase | 2 | 2 | 2 | 2 |
| Uncharacterized | 1 | 1 | 1 | 1 |
| Gliotactin | 1 | 1 | 1 | 1 |
| Glutactin | 5 | 5 | 3 | 7 |
| Uncharacterized | 2 | 1 | 2 | 3 |
| Neurotactin | 2 | 2 | 2 | 2 |
| Uncharacterized H | 1 | 1 | 1 | 1 |
| Neuroglin | 4 | 5 | 4 | 4 |
| **Total** | **36** | **39** | **30** | **45** |
